# Self-Organized Critical Control of Genome Expression: Novel Scenario on Cell-Fate Decision

**DOI:** 10.1101/451260

**Authors:** Masa Tsuchiya, Alessandro Giuliani, Kenichi Yoshikawa

## Abstract

In our current studies on whole genome expression in several biological processes, we have demonstrated the actual existence of self-organized critical control (SOC) of gene expression at both population and single cell level. SOC allows for cell-fate change by critical transition encompassing the entire genome expression that, in turn, is partitioned into distinct response domains (critical states).

In this paper, we go more in depth into the elucidation of SOC control of genome expression focusing on the determination of critical point (CP) and associated distinct critical states in single-cell genome expression. This leads us to the proposal of a potential universal model with genome-engine mechanism for cell-fate change. Our findings suggest that the CP is fixed point in terms of temporal expression variance, where the CP (set of critical genes) becomes active (ON) for cell-fate change (‘super-critical’ in genome-state) or else inactive (OFF) state (‘sub-critical’ in genome-state); this may lead to a novel scenario of the cell-fate control through activating or inactivating CP.

## I. Introduction

A mammalian mature stem cell can reprogram its state to acquire a very different gene expression profile through a few reprogramming stimuli [Takahashi, K., Yamanaka, S., 2016]. This state change (cell-fate change) involves on/off switching on thousands of functionally unique heterogeneous genes in a remarkably coordinated manner [MacArthur, B. D., et al., 2009]. However, there currently exist fundamental difficulties to understand such coordinated control of a large number of genes, in a situation where the lack of a sufficient number of molecules to reach a stable thermodynamic state and the stochastic noise due to the low copy number of specific gene mRNAs per cell provokes a substantial instability of genetic product concentrations which falsifies any gene-by-gene feedback control hypothesis [Raser, J. M., O’Shea, E. K., 2005; Yoshikawa, K., 2002].

In our studies [Tsuchiya, M., et al., 2014-2017; Giuliani, A., et al., 2018], we have demonstrated the existence of self-organization in whole genome expression at both population and single cell level. A fundamental issue is to elucidate the mechanism of the self-organization at the ‘whole genome’ level of gene-expression regulation that is responsible for massive changes in the expression profile through the genome reprogramming inside a small and highly packed cell nucleus.

In order to understand underlying mechanism of self-organization through the massive changes, Per Bak and colleagues [Bak, P., et al., 1987] proposed self-organized criticality (SOC; the Bak-Tang-Wiesenfeld sandpile model) as a general theory of complexity to describe self-organization and emergent order in non-equilibrium systems (thermodynamically open systems), where self-organization is considered to occur at the edge between order and chaos [Langton, C. G., 1990; Kauffman, S. A., 1993], often accompanied by the generation of exotic patterns (good description of SOC in [Jensen, H. J. 1998; Marković, D., Gros, C., 2014]; see current review on criticality in [Muñoz, M. A., 2018]). SOC builds upon the fact that the stochastic perturbations initially propagate locally (sub-critical state), but due to the particularity of the disturbance, the perturbation can spread over the entire system in an autocatalytic manner (super-critical state); as the system approaches its critical point, global collective behavior for self-organization emerges.

The above depicted classical concept of SOC, has been extended to propose a conceptual model of the cell-fate decision (critical-like self-organization or rapid SOC) through the extension of minimalistic models of cellular behavior. The cell-fate decision-making model (a coarse-grained model) considers gene regulatory networks, which adopt an exploratory process, where diverse cell-fate options are first generated by the priming of various transcriptional programs, and then a cell-fate gene module is selectively amplified as the network system approaches a critical state [Halley, J. D., et al., 2009].

We investigated whole genome expression and its dynamics to address the following fundamental questions:

- Is there any underlying principle that self-regulates whole-genome expression?
- Does a universal mechanism exist to guide the self-organization that determines the change in the cell fate?

Our findings suggested that at a specific experimental time point, a transitional behavior of expression profile occurs in the ensemble of genes (e.g., unimodal-bimodal transition in MCF-7 cancer cells [Tsuchiya, M., et al., 2014]). This corresponds to a self-similar behavior around a critical transition point that is evident when expression is sorted and grouped according to temporal variance of expression (normalized root mean square fluctuation: *nrmsf*: **Methods**). On the contrary, randomly shuffled genome expression exhibit Gaussian normal distributions, with no evidence of transitional behavior. This is consistent with the existence of self-organization according to *nrmsf*, that is, *nrmsf* act as an order parameter of self-organization. This suggests that the grouping of expressions averages out expression noise coming from biological and experimental processes allowing to highlight self-organizing behavior through distinct response expression domains (critical states). Therefore, our focus is on averaging behaviors (mean-field approach) of group expression emerged in genome expression.

Our findings of self-organization with critical behavior (criticality) differ from classical and extended SOC models (see **Discussion**) as for the following elements:

i. Occurrence of cell-fate change through erasure of the initial-state sandpile criticality.
ii. Coexistence of critical states (super-critical: high temporal-variance expression; near-critical: intermediate variance expression; sub-critical: low variance expression).
iii. The sub-critical state as a generator of autonomous SOC control (versus non-autonomous classical SOC [Halley, J. D., et al. 2009]), which guides the cell-fate change.
iv. The existence of a potential universal mechanism of cell-fate change over different biological processes.

Two distinct critical behaviors emerged as gene expression values are sorted and grouped: sandpile-type criticality and scaling-divergent behavior (genome avalanche). Sandpile criticality is evident in terms of grouping according to expression fold-change between two different time points, whereas scaling-divergent behavior emerges according to grouping by *nrmsf* (refer to Fig.1 in [Giuliani, A., et al., 2018]). Criticality allows for a perturbation of the self-organization (i.e., change in critical point) due to change in signaling by external or internal stimuli into a cell to have a global impact on the entire genome expression system.

**Figure 1.**
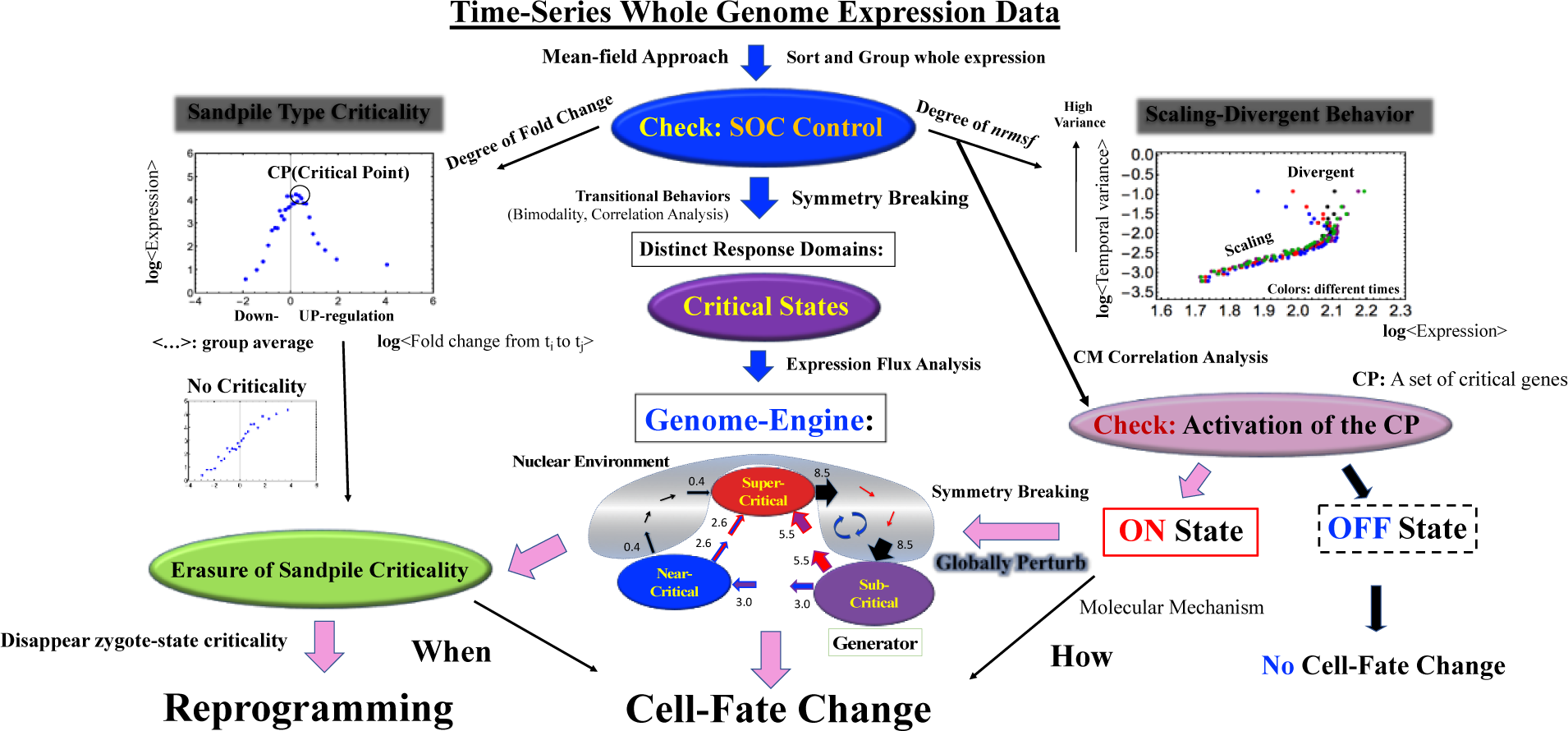
Schematic overview of self-organized critical (SOC) control of whole genome expression. A potential universal model (based on genome-engine hypothesis) for autonomous critical control of whole genome expression is presented (see main text for details).

In this report, we update our previous findings and develop a unified model of cell-fate change as follows:

1. Previously we had some technical difficulties to determine distinct response domains (critical states) for single-cell genome expression in RNA-Seq data, where there are lots of zero-value expression causing specific instability in bimodal transitional behaviors of expression profile. To overcome this drawback, a new correlation metrics based upon the center of mass (CM) of genome expression is developed; this metrics reveals fixed point (regarding temporal expression variance, i.e., time) behavior of critical points (CPs) in both population and single-cell genome expression from embryo development to cell differentiation (see **Section IIA**).
2. These CP behaviors reveal distinct response domains (critical states) in single-cell genome and associated coherent behavior (i.e., CM dynamics) emerged in stochastic expression (stochastic-coherent behavior) in each critical state. These coherent behaviors exhibit a universal genome-engine mechanism [Tsuchiya, M., et al., 2016, 2017] for SOC-control of genome expression (see **Section IIB**).
3. Finally, CP have active or inactive state in population cells, which may suggest a novel cell-fate control mechanism (see overview in **Figure 1;** see more in **Discussion**).

## II. Results

### A. Emergent Genomic Expression System - Self-Organization with Critical Behaviors

#### Critical Behaviors in Temporal Correlation of Genome Expression

To develop a unified view of self-organizing genome expression in distinct biological regulations, the existence of a critical point (CP) plays an essential role in determining distinct response domains (critical states) [Tsuchiya, M., et al., 2016], here we go in depth into specific features of the CP of sandpile type critical behaviors (*sandpile criticality*) (**Figures 2A and 3A**) in genome expression. Our study on HRG-stimulated MCF-7 cancer cells (population level) demonstrated that the temporal group correlation (between groups-correlation) along the order parameter (*nrmsf*) reveals a focal point (FP) when we consider the center of mass (CM) of whole expression (changing in time) as a reference expression point (see Fig.5B in [Tsuchiya, M., et al., 2015]; called ***CM correlation***). The correlation metrics we used to investigate the transition dynamics builds upon the following (very basic) statistical formalization:

1. Genome expression is considered as a *N*-dimensional vector, **G**(*t_i_*) at *t* = *t_i_*: **G**(*t_i_*) can be decomposed into *K* groups of vectors with *n* elements (*N* = *K · n*) according to *nrmsf*: **G**(*t_i_*) = (***g***^1^(*t*_i_), ***g***^2^(*t*_i_), … ***g***^*K*^(*t*_i_)), where ***g***^1^(*t*_i_) and ***g***^*K*^(*t*_i_) are the highest and lowest group vectors of *nrmsf*, respectively and the *k*^th^ group vector with *n* elements is 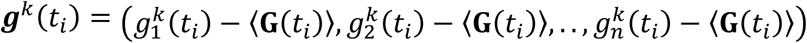 with the average of genome expression, 〈**G**(*t_i_*), i.e., CM of genome expression at *t* = *t_i_* and its vector 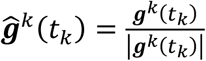. Note that the last group (the lowest *nrmsf*) less than *n* elements is removed from the analysis.
2. Keeping in mind, correlation corresponds to cosine of angle between unit vectors, i.e., inner product of unit vectors (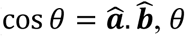: angle; 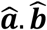: dot product (scalar) of unit vectors: 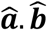). Two different CM correlations are considered:

i. **Spatial CM correlation**: for a fixed time point (*t* = *t_j_*), development of the CM correlation between the first group (highest *nrmsf* group) and other vectors: 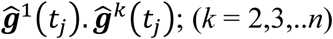. **Figure 2** for HRG-stimulated MCF-7 cells (cell differentiation at population level; **Methods**) shows that the spatial CM correlation exhibits a FP, the intersection of the correlation between different time points (**Figure 2C**). The FP coincides with the CP of sandpile type criticality in terms of *nrmsf* value.
ii. **Temporal CM correlation**: for a fixed group (*k*), development of the CM correlation between the initial and other experimental points: 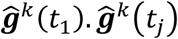 over experimental point, *t_j_*. The temporal CM correlation reveals the CP as divergent behavior (**Figure 2D**) in population cells.

**Figure 2.**
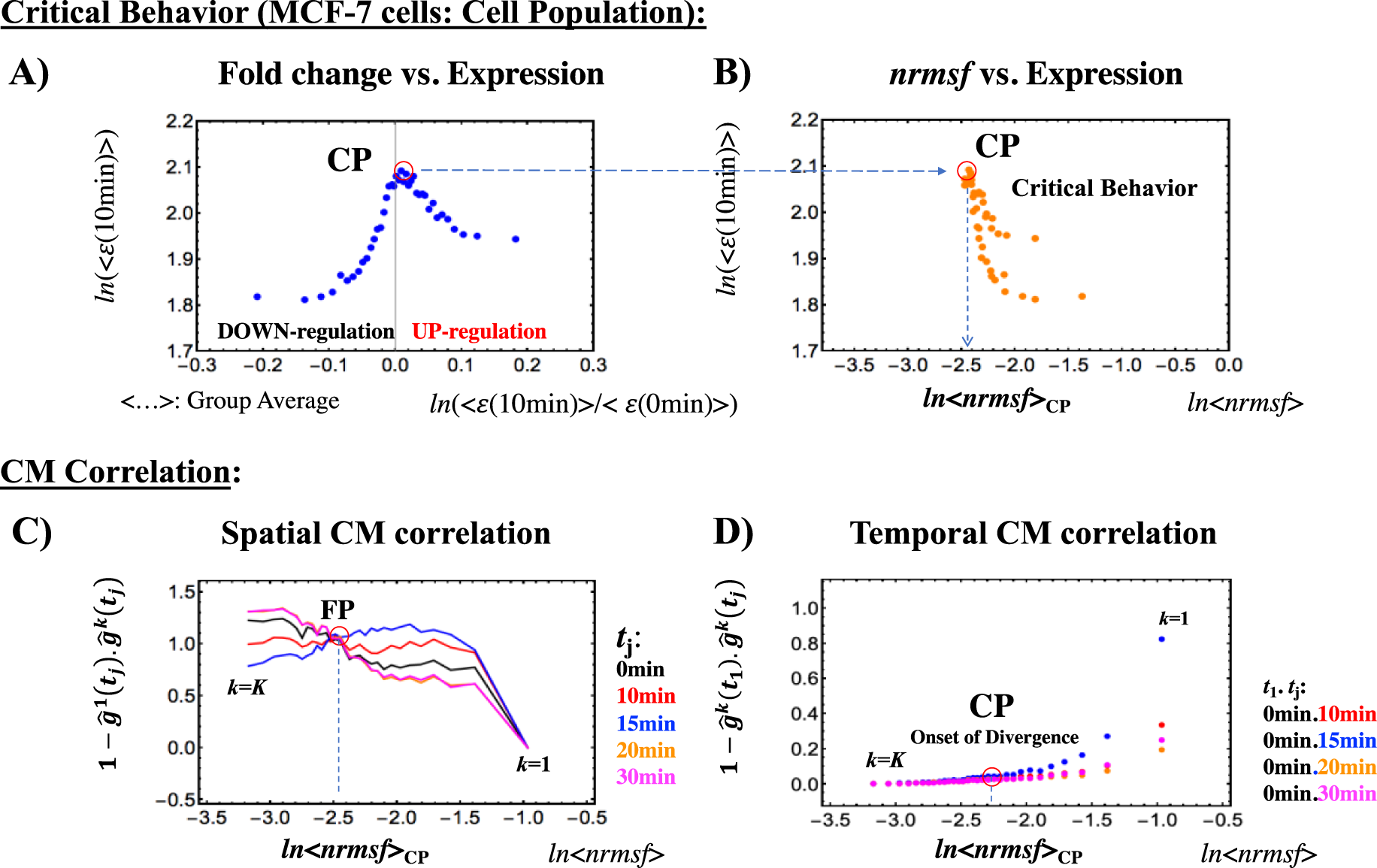
Cell population: Critical behavior with critical point (CP) in HGR stimulated MCF-7 cancer cell differentiation. The first row - whole mRNA expression (microarray data; **Methods**) is sorted and grouped (*K* = 40 groups) according to the fold change in expression (*ε*) between 0min and 10min:

A. Left panel shows sandpile type critical behavior (criticality), where the summit of sandpile corresponds to the CP, *x*- and *y*-axis represents group average value of the fold-change (0-10min) and group average value of expression at 10min, respectively.
B. Right panel shows critical behavior of group average values in *nrmsf* vs. expression plot, where whole expression is sorted and grouped (*K* = 40 groups) according to *nrmsf*. Both *nrmsf* values are the same at the CP (*ln* < *nrmsf* > ∼ -2.5; see **Table 1**). The second row - whole expression is sorted and grouped (*K* = 40 groups) according to *nrmsf*:

C) Left panel shows that 1- spatial CM correlation, 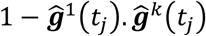 at *t* = *t_j_* exhibits a focal point (FP) around at *ln* <*nrmsf*> ∼ -2.5, which coincides with the CP.
D) Right panel shows that divergent for 1-temporal CM correlation, 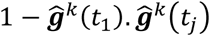 occurs at the CP.

**Table 1.** Activation of the CP on cell-population genome expression associated with the timing of global perturbation. Activation of the CP and Global Perturbation based on the temporal CM correlation and kinetic energy flux analysis, respectively; *Second perturbation for atRA-stimulated HL-60 cells (see more in [Tsuchiya, M., 2016]).

|  | Location of the CP | Activation of the CP | Global Perturbation on the Genome-Engine |
| --- | --- | --- | --- |
| EFG-stimulated MCF-7 cells | $\ln\langle nrmsf \rangle_{CP} \sim -2.5$ | OFF state | None |
| HRG-stimulated MCF-7 cells | $\ln\langle nrmsf \rangle_{CP} \sim -2.5$ | ON state at 15min | 15min - 20min |
| DMSO-stimulated HL-60 cells | $\ln\langle nrmsf \rangle_{CP} \sim -2.5$ | ON state at 12h | 12h - 18h |
| atRA-stimulated HL-60 cells | $\ln\langle nrmsf \rangle_{CP} \sim -2.5$ | ON state at 18h | *12h - 18h |

On the other hand, at single cell level (RNA-Seq data), **Figure 3** shows that whole RNA expression in Th17 cell differentiation (**Methods**) exhibits similar critical behaviors to those of cell population except for critical behaviors in CM correlation of subsequent time points with the initial one (**Figure 3D**).

**Figure 3.**
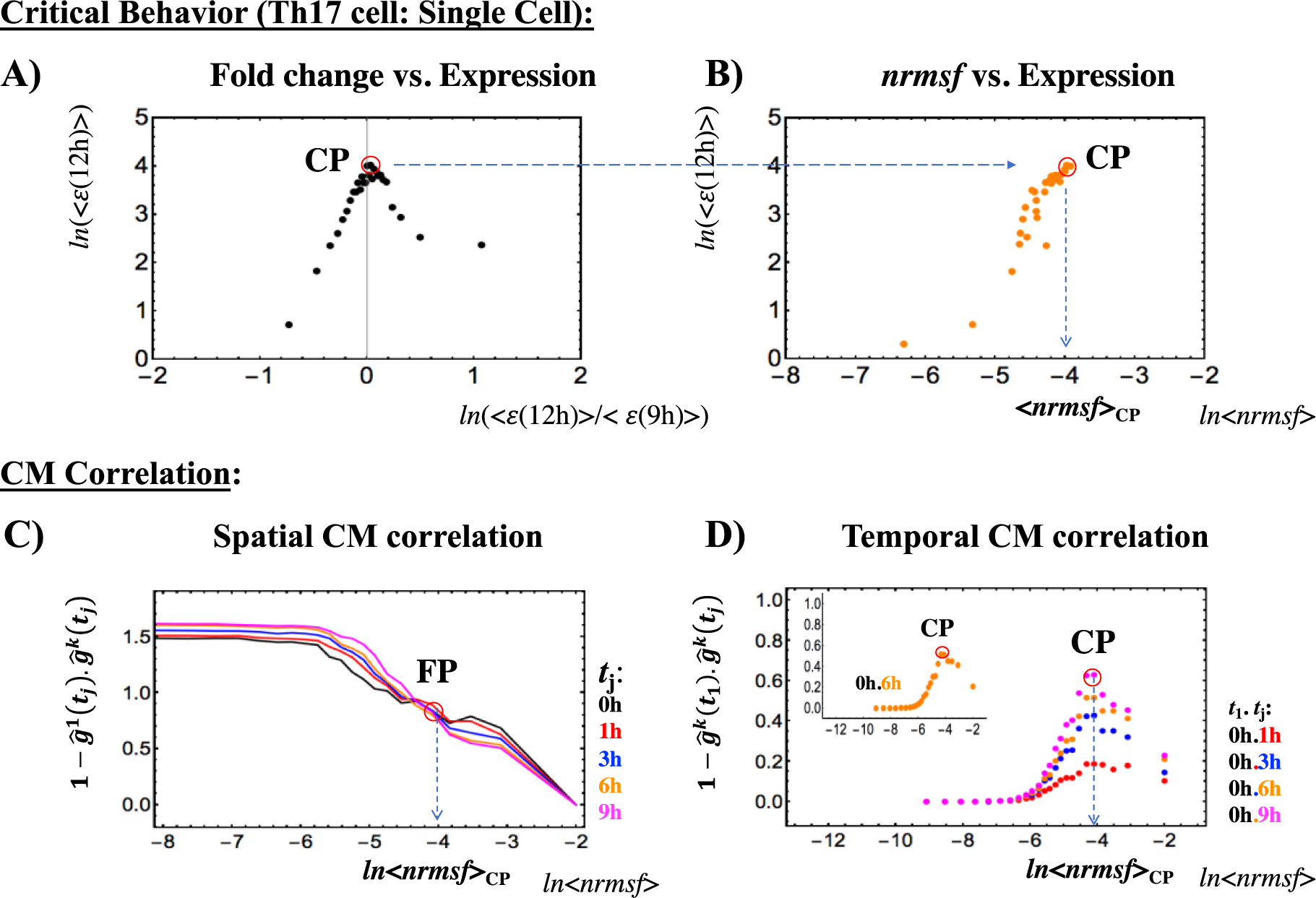
Single cell: Critical behavior with the CP in Th17 immune cell differentiation. Sorting and Grouping of whole RNA expression (RNA-Seq data; Methods) is the same as in **Figure 2**. As shown clearly in **Figure 4**, distinct characteristic difference in criticality from that of population level (**Figure 2**) is in the temporal CM correlation, in which the CP is a point with no differential (vs. divergent point in cell population); this reveals distinct response domains (critical states) in single-cell genome expression (**Figure 4B**).

**Figure 4** confirms two distinguished critical behaviors: 1) Divergent at the CP for micro-array data, 2) Lack of differential (i.e., a critical point) at the CP for single cell RNA-Seq data. Notably both results point to the important fact that the CP is fixed point in time, thus suggesting the existence of a unique critical point in genome expression for a specific biological regulation.

**Figure 4.**
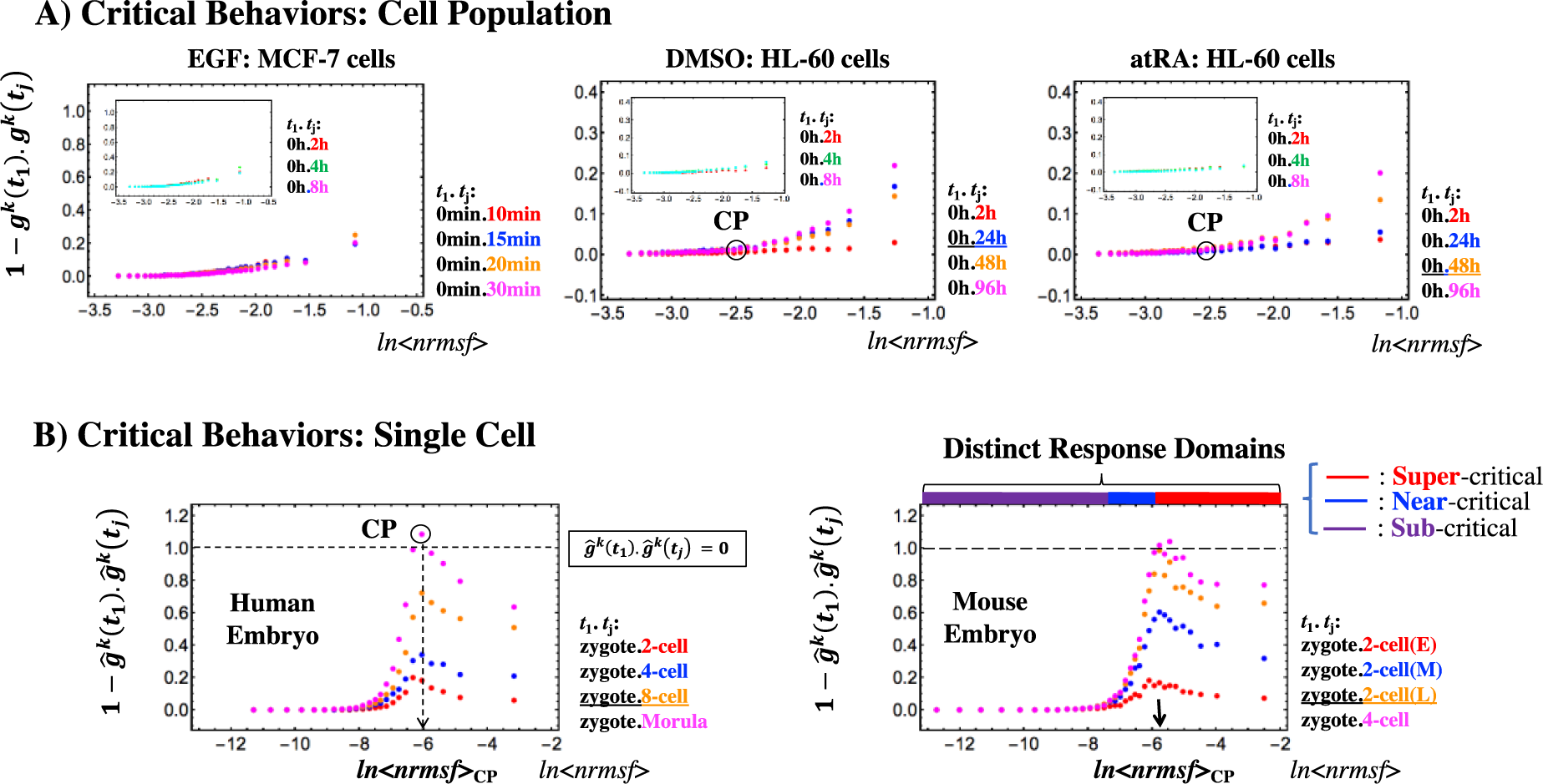
Critical behaviors in the temporal CM correlation: *g*^*K*^(*t*_1_).*g*^*K*^(*t*_*j*_). **A) Cell Population** (Microarray data): Results of 1- temporal CM correlation, (see **Figure 2D**) are shown: left panel: EGF-stimulated MCF-7 breast cancer cells, middle: DMSO-stimulated HL-60 human leukemia cells, and right: atRA-stimulated HL-60 human leukemia cells. EGF-stimulated MCF-7 cells do not show any divergent behavior (compare with **Figures 2D**), which do not go to cell differentiation [Saeki, Y., et al., 2009]. This keep on with time (inset). DMSO- and atRA-stimulated HL-60 cells lead to cell-fate change: at 24h for DMSO and at 48h for atRA [Tsuchiya, M., et al., 2016], where vivid responses in the temporal CM correlation exhibit at 24h (DMSO) and at 48h (atRA), whereas there is almost no divergence before those time points (insets). Both CPs for HL-60 cell exist around *ln*<*nrmsf*> ∼-2.5 (based on sandpile criticality). **B) Single cell** (RNA-Seq data): critical behaviors for 1- temporal CM correlation (see **Figure 3D**) for human (left panel) and mouse (right) embryo development are shown. Mouse and human CPs are fixed point, where the CPs exist around *ln*<*nrmsf*> ∼-6.0 (values confirmed by the analysis based on **Figures 2, 3**; see more in **Table 2**). Notably, both CPs pass zero correlation (random like behavior) after 8-cell state (human) and late 2-cell state (mouse), which coincides with the timing of reprogramming - time for the erasure of initial-state sandpile criticality [Tsuchiya, M., et al., 2016, 2017]. Distinct response domains are shown: super-critical: red line; near-critical: blue; sub-critical: purple (refer to **Table 2**).

**Table 2.** Critical states of single-cell genome expression with critical point (CP).

| Mouse Embryo Development | Human Embryo Development | Th17 cell differentiation |
| --- | --- | --- |
| **Critical States:** | **Critical States:** | **Critical States:** |
| **Super-critical:** 4203 RNAs<br>$-5.8 < \ln\langle nrmsf \rangle$ | **Super-critical:** 3088 RNAs<br>$-6.1 < \ln\langle nrmsf \rangle$ | **Super-critical:** 2373 RNAs<br>$-4.09 < \ln\langle nrmsf \rangle$ |
| **Near-critical:** 5302 RNAs<br>$-7.5 < \ln\langle nrmsf \rangle < -5.8$ | **Near-critical:** 6589 RNAs<br>$-8.1 < \ln\langle nrmsf \rangle < -6.1$ | **Near-critical:** 6190 RNAs<br>$-6.3 < \ln\langle nrmsf \rangle < -4.1$ |
| **Sub-critical:** 7636 RNAs<br>$\ln\langle nrmsf \rangle < -7.5$ | **Sub-critical:** 6982 RNAs<br>$\ln\langle nrmsf \rangle < -8.1$ | **Sub-critical:** 4571 RNAs<br>$\ln\langle nrmsf \rangle < -6.3$ |
| **Critical Point:** | **Critical Point:** | **Critical Point:** |
| $\ln\langle nrmsf \rangle_{CP} \sim -5.8$<br>**CM correlation of the CP:**<br>**Zero**<br>**After 8-cell state** | $\ln\langle nrmsf \rangle_{CP} \sim -6.1$<br>**CM correlation of the CP:**<br>**Zero**<br>**After Late 2-cell state** | $\ln\langle nrmsf \rangle_{CP} \sim -4.1$<br>**CM correlation of the CP:**<br>**Never Pass Zero** |

Dynamics of the CM correlation reveals additional features of the CP that

i. EGF-stimulated MCF-7 cells, where no cell-differentiation occurs [Saeki, Y., et al., 2009]], does not show any divergent behavior (**Figure 4A**; confirmed further by repetitive data; **Methods**). Sand-pile criticality is present in EGF-stimulated MCF-7 too, but no characteristic change of criticality such as the erasure of initial-state criticality occurs in time course of EGF stimulation (i.e., no global change in genome expression) [Tsuchiya, M., et al., 2016]. This suggests that **the CP possesses active or inactive state**, i.e., ON or OFF expression state for set of genes (**critical gene set**) corresponding to the CP; in EGF-stimulated MCF-7 cells, the CP is in the inactive (OFF mode) state. DMSO- and atRA-stimulated HL-60 cells further support this condition of the CP in that before the cell fate change (24h: DMSO;48h: atRA; see **Table 1**), there are no divergent behaviors in the temporal CM correlation (insets of **Figure 5A**). **This finding suggests that activation of critical gene set (CP) plays an important role in cell-fate change.** Our studies revealed the occurrence of global perturbation on the self-organization before the genome-state change [Tsuchiya, M., et al., 2016]. It is intriguing to observe that the timing of activation of the CP coincided with that of the global perturbation (**Table 1**); at the beginning of the global perturbation, the activation of the CP occurs (12h in DMSO-stimulated HL-60 cells and 15min in HRG-stimulated MCF-7 cells) (Fig. 14 and see also no global perturbation in EGF-stimulated MCF-7 cells in [Tsuchiya, M., et al., 2016]). This supports the genome-engine mechanism (**Section IIB**) that implies the activation of the CP has an impact on the entire genome expression provoking a generalized perturbation. Therefore, **the elucidation of underlying mechanism, which guides the state (activation or inactivation) of the CP, is expected to predict when and how a cell-fate change occurs, i.e., a novel cell-fate control mechanism for cancers, iPS cells, stem cells and so forth**(see **Discussion**).
ii. In the case of embryo development, the temporal CM correlation for the CP traverses zero value (correspondent to random-like behavior) after the reprogramming event (after late 2-cell and 8-cell state for mouse and for human, respectively) (**Figure 4B**), which coincides with the erasure of the sandpile-type critical point (CP) (refer to Fig.2 in [Giuliani, A., et al., 2018]). In biological terms this corresponds to the erasure of the initial stage of embryogenesis (driven by maternal heredity); on the contrary, the CP for Th17 cell differentiation does not pass zero correlation (**Figure 3D**). Thus, **additional condition for the embryonic reprogramming is suggested whether or not the temporal CM correlation pass zero,** and
iii. The results imply the existence of cell-type specific fixed-point CP, but we need further investigation on time-series genome expression data for different biological systems to confirm this conjecture.

**Figure 5.**
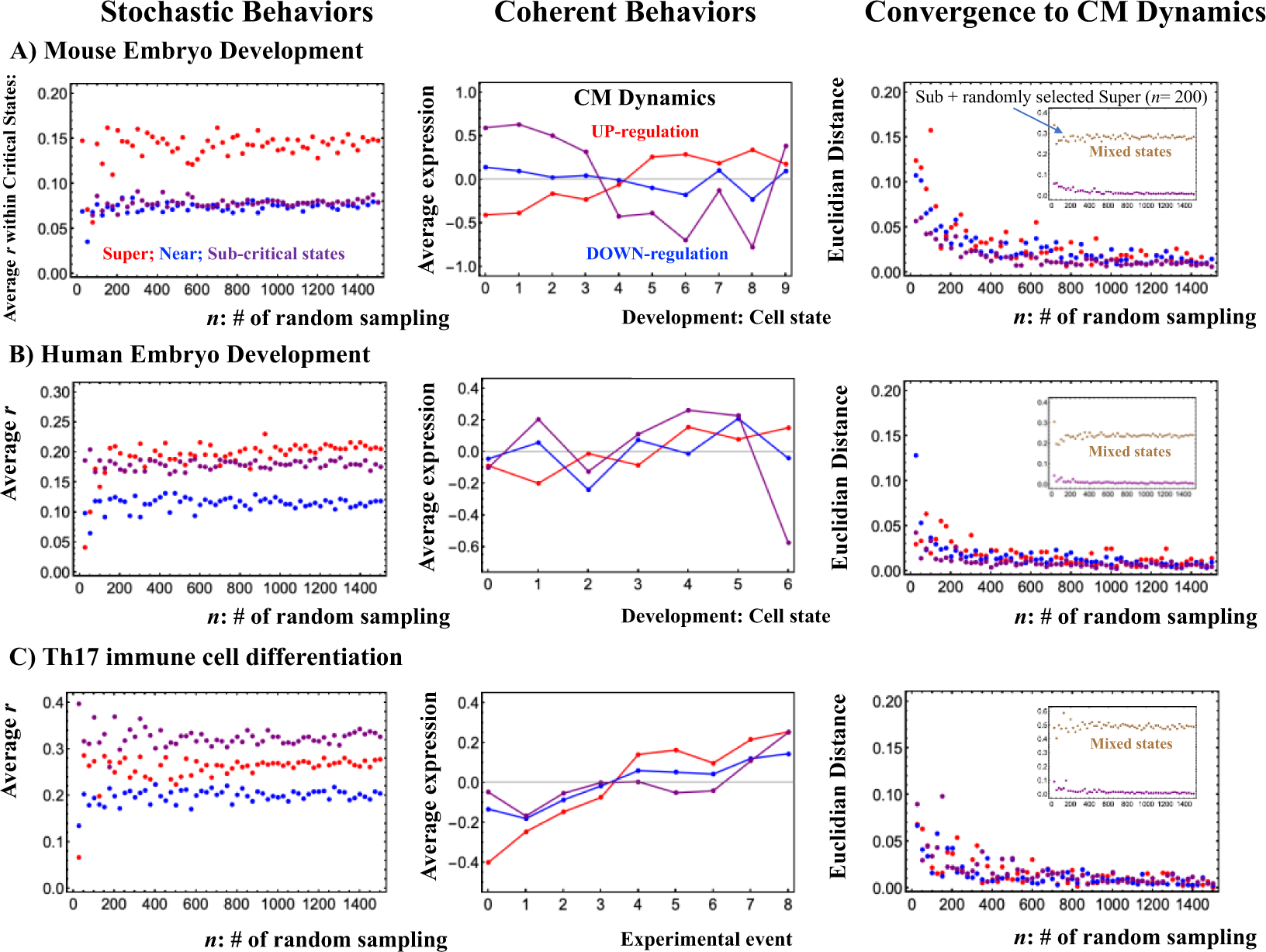
Stochastic-coherent behaviors in single-cell genome expression. A) mouse embryo development; B) human embryo development; C) Th17 immune cell differentiation. **Left panel**: Average Person correlation (200 repetitions) between randomly selected expression within a critical state (red: super-; blue: near-; purple: sub-critical state) shows stochastic expression. **Middle panel**: CM dynamics - time development of the average expression of each critical state (development in cell state as time development) is shown. Each integer point in the *x* axis represents development of cell state for embryo development and experimental time for Th17 cell differentiation (refer to Methods for corresponding cell state and experimental time). **Right panel**: Convergence (by Euclidian distance between dynamics of the CM and average expression of randomly selected expression) to the CM of randomly selected expression from each critical state shows law of large number in each critical state, but no longer this is true in addition of randomly selected super-critical expression (*n* = 200) into sub-critical expression (inset).

It is important to note that groups of low-*nrmsf* presenting elevated CM correlations in time do not point to a no-response situation; on the contrary they behave in a highly coherent manner to generate autonomous SOC mechanism (see **Section IIB**).

#### Determination of Critical States in Single Cell Genome Expression

In our previous studies on microarray data (at cell population level), we checked the transition of expression profile by means of Sarle’s bimodality coefficient putting in evidence of distinct response domains (super-, near- and sub-critical domains according to temporal variance of expression) were evident in genome expression [Tsuchiya, M., et al., 2016]. In the case of single-cell studies, we met some difficulties in determining the response domains. Gene expression in the single cell RNA-Seq data has many zero values: this fact cannot be considered as a mere artifact stemming from technical problems, but rather a consequence of the “toggle-switch” mechanism of gene regulation [Kim, K.Y., Wang, J., 2007] - essential for understanding regulation mechanisms. However, these zero values create instability in the assessment of bimodality for the determination of distinct response domains (critical states) as well as problems of divergence (logarithm of zeros) in the scaling analysis in log-log plots. We overcame this problem by a bootstrap approach: instead of computing a single group correlation we consider the average of between-groups correlation relative to 200 repetitions of randomly selected expressions with a critical state. The existence of a threshold *n* at around 50 randomly picked genes [Censi, F., et al., 2011; Tsuchiya, M., et al., 2015], which allows to reproduce the correlation properties of the entire genome with a random choice of *N* genes with *N* > *n* is a further proof of the reliability of our strategy such as the CM correlation analysis in this report.

Genome expression at single cell level presents the same sand-pile type criticality observed at cell population level [Tsuchiya, M., et al., 2015, 2016]. We investigate the suitability of CM correlation as metrics whether to capture critical behavior in genome expression; **Figure 4B** clearly shows that the temporal CM correlation manifest distinct response domains according to *nrmsf*: low-variance expression (sub-critical state) for region of flatten correlation (region of perfect correlation), intermediate-variance for near-critical state from the edge of flatten correlation to the CP, and high-variance expression for super-critical state above the CP. These critical states possess coherent-stochastic behaviors - coherent behavior emerged from ensemble of stochastic expression [Tsuchiya, M., et al., 2016] (**Figure 5**). Importantly, this suggests that self-organizing principle of genome expression is the SOC control of overall expression in single-cell as well as in cell population.

Note that Pearson correlation, i.e., reference point as time average of expression for each gene does not shows the critical point (data not shown); this points to the necessity of considering integrated genome expression (and not chasing for single gene markers) in order to get rid of global cell response.

Next, we examine whether or not a universal mechanism of autonomous SOC control of genome expression - genome engine mechanism exists in single cell genome (see genome engine mechanism for cell population in [Tsuchiya, M., et al., 2016]).

### B. Genome Engine Hypothesis - A Potential Universal Mechanism of Self-Organized Critical Control of Genome Expression

Single-cell genome expression shows distinct critical states with stochastic-coherent behaviors (**Figure 5**) - distinct coherent behavior (CM) emerged from stochastic expression, where stochastic expression in each critical state follow law of large numbers, but mixing state does not converge to the CM of critical state due to distinguished coherent dynamics of critical states. The stochastic-coherent behaviors, which discloses that the CM of critical state represents its dynamics, are exactly same as ones in cell population (Fig. 10 in [Tsuchiya, M., et al., 2016]) and thus, dynamic expression flux analysis between critical states can apply to reveal the genome-engine mechanism in single cell (below) for description of how autonomous SOC control of genome expression occurs.

**Figure 6A** shows that the sub-critical state acts as internal source of expression flux and super-critical state as sink. Sub-super cyclic flux forms a dominant flux flow, which in turn generates strong coupling between the super- and sub-critical states accompanied by their anti-phase expression dynamics [Tsuchiya, M., et al., 2016], makes its change in oscillatory feedback, and thus sustains autonomous SOC control of overall gene expression. The formation of a dominant cyclic flux provides a universal genome-engine metaphor of SOC control mechanisms pointing to a universal mechanism in gene-expression regulation of mammalian cells for both population and single cell levels. Kinetic energy flux (**Figure 6B**; **Methods**) in single cell clearly shows that global perturbation occurs on the genome-engine (i.e., self-organization of whole genome expression), where its timing coincides with that of the cell-fate change (refer to Fig. 2 in Giuliani, A., et al., [2018]). Hence, this suggests that in regard to embryonic reprogramming, the activation of the CP points to when the temporal CM correlation of the CP from initial state (zygote) passes zero-correlation (i.e., erasure of initial-state CP memory).

**Figure 6.**
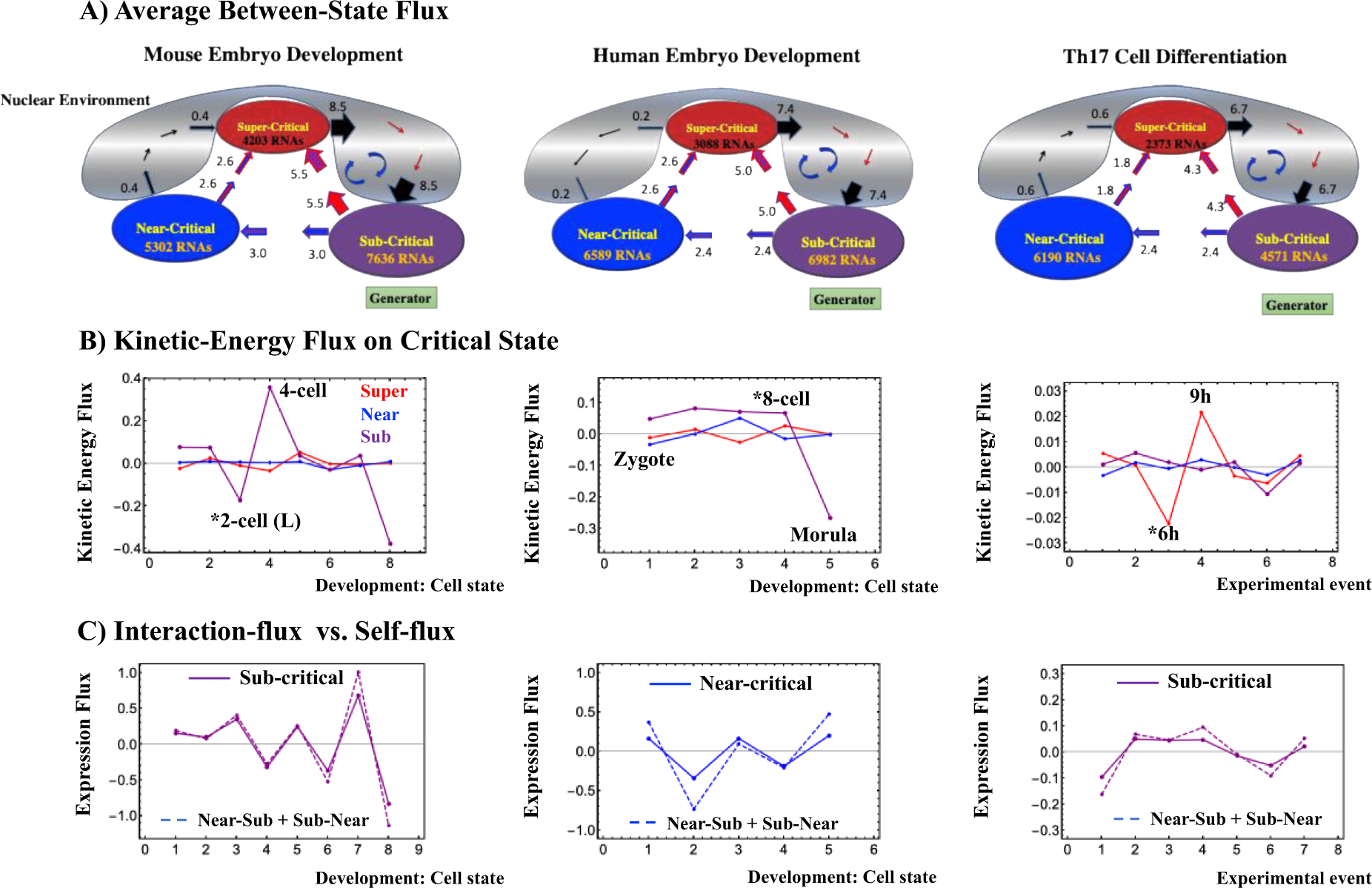
Genome engine mechanism through average between-state flux. A) Sub-super cyclic flux forms a dominant flux flow to establish genome engine mechanism: the sub-critical state acts as a “large piston” for short moves (low-variance expression) and the super-critical state as a “small piston” for large moves (high-variance expression) with an “ignition switch” (near-critical state with a critical point) are connected through a dominant cyclic state flux as a “camshaft”, resulting in the anti-phase dynamics of two pistons. B) Kinetic energy flux (**Methods**) shows that the timing (indicated by asterisk) of global perturbation (involved in more than one critical state) coincides with that of cell-fate change. C) Interaction flux (dashed lines) between Near-Sub and Sub-Near determines the net flux (self-flux) of sub-critical states (mouse embryo: left panel and Th17 cell: right panel) and near-critical state (human embryo: middle panel). This shows that activation of the CP affects the entire genome through the Sub-Near interaction flux (see main text).

Furthermore, the genome-engine in control of dynamics of whole genome expression (**Figure 6A**) demonstrates how perturbation on the sub-critical state (generator of SOC-control) affects the entire genomic system, which shows that the activation of the CP (the edge of the near-critical state) can induce a global impact on the entire genome expression through interaction between near- and sub-critical states (Fig.13 for cell population and Discussion in [Tsuchiya, M., et al., 2016]; **Figure 6C** for single cell).

## III. Discussion

We further demonstrated for several distinct biological regulations from embryo development to cell differentiation that self-organized criticality (SOC) control of genome expression provides a potential universal model (**Figure 1**) of self-organization as the genome-engine mechanism, where a highly coherent behavior of low-variance genes (sub-critical state) generates a dominant cyclic expression flux with high-variance genes (super-critical state) to develop autonomous critical control system. This explains the coexistence of critical states (distinct expression response domains) through critical point (CP); *nrmsf* (temporal expression variance) acting as the order parameter for the self-organization. The genome-engine mechanism rationalizes how the change in criticality affect the entire genome expression. This model explains how collective behavior of stochastic low-variance expression (stochastic-coherent behavior) guides reprogramming event in early mouse embryo development. A highly coherent behavior emerges from stochastic low-variance expression (sub-critical state) and acts as the engine generator, driving the genome through a critical transition state right after the late 2-cell state [Tsuchiya, M., et al., 2017]. Recent study shows that the dynamics of high order structure of chromatin exhibits liquid like behavior [Maeshima, K., et al., 2016], which could be crucial characteristic to enable the genome to conduct SOC gene expression control for cell-fate determination.

Another relevant finding is the existence of a fixed critical gene set corresponding to the CP for specific biological regulation and the activation of these genes (CP) is essential for the occurrence of cell-fate change; conversely inactivation of the CP does not go to cell-fate change. Furthermore, the genome-engine suggests that activation mechanism of the CP should elucidate how the global perturbation occurs on self-organization through change in signaling by external or internal stimuli into a cell. This, at least in principle, could be the key to understand when and how the cell-fate change occurs.

The activation and inactivation of the CP suggests that there may be another layer of a macro-state (genome state) composed of distinct micro-critical states (found by us). The activation of the CP makes the genome-state to be a ‘super-critical’ to guide the cell-fate change - super-critical after activation of the CP: HRG-stimulated MCF-7 cells, and DMSO- and atRA-stimulated HL-60 cells, whereas before the activation of their CPs, the genome states are ‘sub-critical’, and the genome state of EGF-stimulated MCF-7 cells remains ‘sub-critical’ (no cell-fate change) all the way (**Figure 4A**).

Regarding reprogramming of human and mouse embryo cells (single cells), our result suggests that reprogramming occurs after the memory of initial state of the CP (e.g., zygote state) is lost (i.e., after the temporal CM correlation of the CP group pass zero-correlation: **Figure 4B**); the timing of this correlation event coincided with the timing of erasure of initial-state criticality for mouse and human embryo developments [Tsuchiya, M., et al., 2017; Giuliani, A., et al., 2018]. Furthermore, the timing of global perturbation on single-cell genome expression coincides with that of reprogramming (**Figure 6B**), which suggests that the activation of the CP occur when the initial-state CP memory is erased.

Further studies on these matters are needed to clarify the underlying fundamental mechanism, and the development of a theoretical foundation for the autonomous critical control mechanism in genome expression as revealed in our findings is expected to open new doors for a universal control mechanism of the cell-fate change.

As for now we can safely affirm the strong interaction among genes with very different expression variance and physiological role push for a complete re-shaping of the current molecular-reductionist view of biological regulation looking for single ‘significantly affected’ genes for the explanation of regulation processes. The view of the genome acting as an integrated dynamical system is here to stay.

## IV. Methods

### Biological Data Sets

We analyzed mammalian transcriptome experimental data for seven distinct cell fates in different tissues:

#### Cell population

1. Microarray data of the activation of ErbB receptor ligands in human breast cancer MCF-7 cells by EGF and HRG; Gene Expression Omnibus (GEO) ID: GSE13009 (N = 22277 mRNAs; experimental details in [Saeki Y, et al., 2009]), which has 18 time points: t_1_ = 0, t_2_ = 10,15, 20, 30, 45, 60, 90min, 2, 3, 4, 6, 8, 12, 24, 36, 48, t_T = 18_ = 72h. Two replicas (rep 1 and rep 2) of the same experiment were taken into consideration.
2. Microarray data of the induction of terminal differentiation in human leukemia HL-60 cells by DMSO and atRA; GEO ID: GSE14500 (N = 12625 mRNAs; details in [Huang, S., et al., 2005]), which has 13 time points: t_1_ = 0, t_2_ = 2, 4, 8, 12, 18, 24, 48, 72, 96, 120, 144, t_T=13_ =168h.

#### Single cell

3. RNA-Seq data of early embryonic development in human and mouse developmental stages in RPKM values; GEO ID: GSE36552 (human: N = 20286 RNAs) and GSE45719 (mouse: N = 22957 RNAs) (experimental details in [Yan, L., et al., 2013]) and [Deng, Q., et al., 2014], respectively). We analyzed 7 human and 10 mouse embryonic developmental stages: Human: oocyte (*m* = 3), zygote (*m* = 3), 2-cell (*m* = 6), 4-cell (*m* = 12), 8-cell (*m* = 20), morula (*m* = 16) and blastocyst (*m* = 30), Mouse: zygote (*m* = 4), early 2-cell (*m* = 8), middle 2-cell (*m* = 12), late 2-cell (*m* = 10), 4-cell (*m* = 14), 8-cell (*m* = 28), morula (*m* = 50), early blastocyst (*m* = 43), middle blastocyst (*m* = 60) and late blastocyst (*m* = 30), where *m* is the total number of single cells.
4. RNA-Seq data of T helper 17 (Th17) cell differentiation from mouse naive CD4+ T cells in RPKM (Reads Per Kilobase Mapped) values, where Th17 cells are cultured with anti-IL-4, anti-IFNγ, IL-6 and TGF-β, (details in [Ciofani, M., et al., 2012]; GEO ID: GSE40918 (mouse: N = 22281 RNAs), which has 9 time points: t_1_ = 0, t_2_ = 1, 3, 6, 9, 12, 16, 24, t_T_=9 = 48h. For each time point, sample numbers are as follows: GSM1004869-SL2653 (t= 0h); GSM1004941-SL1851 (t = 1h); GSM1004943-SL1852 (t = 3h); GSM1005002-SL1853 (t= 6h); GSM1005003-SL1854 (t= 9h); GSM1004934-SL1855 (t= 12h); GSM1004935,6,7-SL1856, SL8353, SL8355 (t= 16h; average of three data); GSM1004942-SL1857 (t = 24h); GSM1004960-SL1858 (t= 48h).

For microarray data, the Robust Multichip Average (RMA) was used to normalize expression data for further background adjustment and to reduce false positives [Bolstad, B. M., et al., 2003; Irizarry, R. A., et al, 2003; McClintick, J. N., Edenberg, H. J., 2006].

For RNA-Seq data, RNAs with RPKM values of zero over all of the cell states were excluded. Random real numbers in the interval [0, *a*] generated from a uniform distribution were added to all expression values in the analysis of sandpile criticality (**Figures 2A, 3A**; no addition of random numbers in the rest of the figures). This procedure avoids the divergence of zero values in the logarithm. The robust sandpile-type criticality through the grouping of expression was checked by changing a positive constant: *a* (0 < *a* < 10); we set *a* = 0.01. Note: The addition of large random noise (*a* ≫ 10) destroys the sandpile CP.

### Normalized Root Mean Square Fluctuation (nrmsf)

*Nrmsf* (see more Methods in [Tsuchiya, M., et al., 2015]) is defined by dividing *rmsf* (root mean square fluctuation) by the maximum of overall {*rmsf*_i_}:

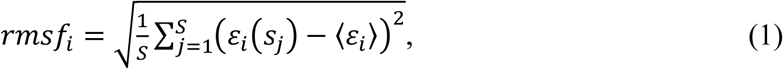

where *rmsfi* is the *rmsf* value of the *i^th^* RNA expression, which is expressed as *ε_i_*(*s_j_*) at a specific cell state *s_j_* or experimental time (e.g., in mouse embryo development, *S* = 10: *s*_1_ = zygote, early 2-cell, middle 2-cell, late 2-cell, 4-cell, 8-cell, morula, early blastocyst, middle blastocyst and *s*_10_ = late blastocyst), and 〈*ε_i_*〉 is its expression average over the number of cell states. Note: *nrmsf* is a time-independent variable.

### Single Cell Expression Flux Dynamics

Here, to describe genome engine mechanism on single cell genome expression, the key fact is that dynamics of coherent behavior emerged from stochastic expression in distinct critical states (coherent-stochastic behavior: CSB) follows the dynamics of the CM of a critical state (**Figure 6**). We have developed expression flux approach to reveal dynamic interaction flux between critical states [Tsuchiya, M., et al., 2016-2018]. The CSB in a critical state corresponds to the scalar dynamics of its CM, *X*(*s*_j_), where *X*(*s*_j_) represents the numerical value of a specific critical state (super-, near- or sub-critical state). The expression flux between critical states is interpreted as a non-equilibrium system and evaluated in terms of a dynamic network of effective forces, where interaction flux is driven by effective forces between different critical states and can be described by a second-order time difference. It is important to note that the oscillatory phenomenon interpreted using a second-order time difference equation with a single variable is equivalent to inhibitor-activator dynamics given by a couple of first-order time difference equations with two variables. Flux dynamics approach is further developed to analyze quantitative evaluation of the degree of non-harmonicity and time reversal symmetry breaking in nonlinear coupled oscillator systems [Tsuchiya, M., et al., 2018].

Basic formulas of expression flux dynamics are given as follows:

#### Net self-flux of a critical state

The net self-flux: IN flux - OUT flux, the effective force on a critical state, represents a positive sign for incoming force (net IN self-flux) and a negative sign for outgoing force (net OUT self-flux); the CM from its average over all cell states represents up- (down-) regulated expression for the corresponding net IN (OUT) flux.

The effective force is a combination of incoming flux from the past to the present and outgoing flux from the present to the future cell state:

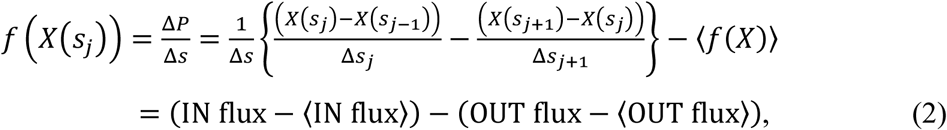

where Δ*P* is the change in momentum with a unit mass (i.e., the impulse: *F*Δ*s* = Δ*P*) and natural log of average (<…>) of a critical state, 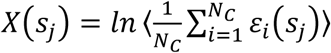 with the *i*^th^ expression *ε_i_*(*S_j_*) at the *j*^th^ cell state, *s* = *s*_j_ (*N_C_* = the number of RNAs in a critical state; the average of net self-flux over the number of critical states, <*f*(X)> = <INflux> - <OUTflux>. Here, scaling and critical behaviors occur in log-log plots of group expression, where natural log of average value associated with group expression such as *ln*<*nrmsf*> and *ln* <*expression*> is taken; thus, in expression flux, we consider natural log of average expression (CM) of a critical state.

It is important to note that each embryo state in terms of number of cells is considered as a statistical event (note: a statistical event does not necessarily coincide with a biological event) and its development as time arrow (time-development) when evaluating average of group expression, fold change in expression and temporal expression variance (*nrmsf*). This implies a time interval in the dynamical system (Equation (2)) is evaluated as difference in event, i.e., Δ*s*_j_ = *s*_j_ - *s*_j-1_ = 1 and Δ *s* = *s*_j+1_ - *s*_j-1_ = 2, as well as difference in experimental times such as in cell differentiation (note: actual time difference can be considered as scaling in time). Then, we evaluate force-like action in expression flux.

#### The interaction flux of a critical state

The interaction flux represents flux of a critical state *X*(*s*_j_) with respect to another critical state (Super, Near, Sub) or the environment (E: milieu) *Y_j_* can be defined as:

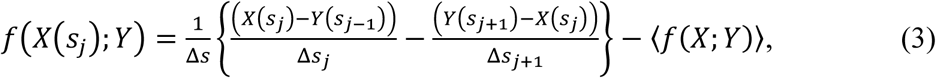

where, again, the first and second terms represent IN flux and OUT flux, respectively, and the net value, IN flux- OUT flux, represents incoming (IN) interaction flux from *Y* for a positive sign and outgoing (OUT) interaction flux to *Y* for a negative sign. *Y* represents the numerical value of a specific critical state or the environment, where a state represented by *Y* is deferent from one by *X.*

With regard to global perturbation event, the net kinetic energy flux [Tsuchiya, M., et al., 2016] clearly reveals it in a critical state (**Figure 6C**):

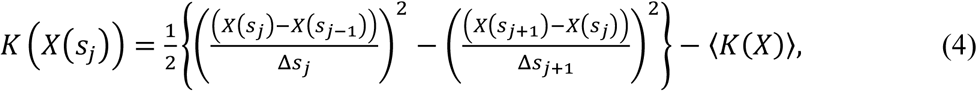

where the kinetic energy of the CM for the critical state with unit mass at *s* = *s_j_* is defined as ½. *ν*(*s_j_*)^2^ with average velocity: 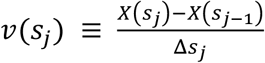.

#### Net self-flux as summation of interaction fluxes

Due to the law of force, the net self-flux of a critical state is the sum of the interaction fluxes with other critical states and the environment:

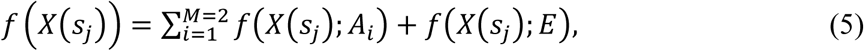

where state *A_i_* ∈ {Super, Near, Sub} with *A_i_* ≠ *X*, and *M* is the number of internal interactions (*M* = 2), i.e., for a given critical state, there are two internal interactions with other critical states. Equation (5) tells us that the sign of the difference between the net self-flux and the overall contribution from internal critical states, 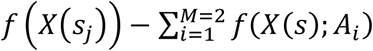, reveals incoming flux (positive) from the environment to a critical state or outgoing flux (negative) from a critical state to the environment.

Here, we need to address the previous result of expression flux dynamics in mouse single-cell genome expression [Tsuchiya, M., et al., 2017], where expression of a critical state is taken as 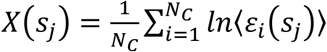, which has different ordering of operations: first taking natural log of expression and then, average operation. Hence, in flux dynamics, we examine whether or not mathematical operation between averaging and natural log, i.e., operation between 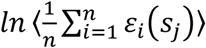 and 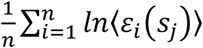 can be exchanged (mathematically commuted). In microarray data, flux behaviors do not change much between these action ordering (almost the same: commuted). Whereas in RNA-Seq data, they are not commuted due to its data structure with lots of zero values; adding small random noise into log of expression, *ln*〈*ε_i_*(*S_j_*)〉 (previous result) makes good effect (noise-sensitive), but not in 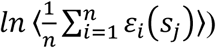: noise-insensitive (this report). Although detail dynamics of interaction flux changes by taking different action-orderings in RNA-Seq data (e.g., Fig. 6 in [Tsuchiya, M., et al., 2017]), two important characteristics in genome-engine: the formation of dominant cyclic flux between super- and sub-critical states and generator role of the sub-critical state do not change (invariant features); thus, we conclude that concept of the genome-engine is robust.

## Contributions

MT initiated the project; MT, AG and KY designed the study; MT developed the study and analyzed data; AG and KY provided theoretical support; MT, AG and KY wrote the manuscript.

## Acknowledgments

MT sincerely thanks the SEIKO Life Science Laboratory, Osaka, Japan and his family for allowing him to complete this research project.

